# Disease responses of hexaploid spring wheat (*Triticum aestivum*) culms exhibiting premature senescence (dead heads) associated with *Fusarium pseudograminearum* crown rot

**DOI:** 10.1101/2020.05.13.094979

**Authors:** N. L. Knight, B. Macdonald, C. Percy, M. W. Sutherland

**Affiliations:** Centre for Crop Health, Faculty of Health, Engineering and Sciences, University of Southern Queensland, Toowoomba, Queensland, 4350, Australia; Queensland Department of Agriculture and Fisheries, Leslie Research Facility, Toowoomba, Queensland, 4350, Australia

**Keywords:** field experiment, *Fusarium*, microscopy, quantitative PCR, time-course

## Abstract

Hexaploid spring wheat (*Triticum aestivum*) may exhibit significant crown rot disease responses to infection by *Fusarium pseudograminearum*, with a range of susceptibility levels available in commercial cultivars. Dry conditions during grain-fill may lead to the expression of prematurely senescing culms, which typically fail to set grain. Assessment of hexaploid spring wheat plants exhibiting both non-senescent and prematurely senescent culms was performed using visual discolouration, *Fusarium pseudograminearum* biomass, vascular colonisation and quantification of wheat DNA in culm sections sampled at three different heights above the crown and at the peduncle. A comparison of these parameters at four time points from milk development, when senescent culms are first observed, to maturity was conducted. Samples from six commercial cultivars were collected in 2014 from Narrabri and Tamworth, New South Wales and Wellcamp, Queensland. Prematurely senescent culms exhibited greater visual discolouration, *Fusarium pseudograminearum* biomass and vascular colonisation than non-senescent culms in each cultivar. Colonisation of xylem and phloem tissue was extensive in the basal portions of prematurely senescent culms (36 to 99%), and suggests significant impacts on water and nutrient movement during crown rot disease. Maturation coincided with significant changes in *Fusarium pseudograminearum* biomass and vascular colonisation. Wheat DNA content varied among cultivars, culm conditions, culm sections and sampling times. The variation in the severity of disease states between culms of the same plant suggests that the timing of initiation of infection in individual culms may vary.

## Introduction

*Fusarium* crown rot is a significant disease of wheat around the globe (Burgess *et al.*, 2001; Kazan & Gardiner, 2018). In Australia, crown rot of wheat causes an estimated $79 million per annum in yield losses (Murray & Brennan, 2009), with significant losses of up to 35% also reported in the Pacific North West of the United States (Smiley *et al.*, 2005). A review by Alahmad *et al.* (2018) outlines the current understanding of crown rot in Australia, where the major causal agent is the fungus *Fusarium pseudograminearum*, with *F. culmorum* also contributing significantly in several growing regions with cooler, moister conditions (Backhouse *et al.*, 2004). *F. pseudograminearum* has also been reported as an important crown rot pathogen in China, Europe, North Africa, North America, South Africa and West Asia (Agusti-Brisach *et al.*, 2018; Burgess *et al.*, 2001; Kazan & Gardiner, 2018).

Crown rot symptoms in cereals are described as honey brown lesions occurring on leaf sheaths, internodes and nodal tissues (Burgess *et al.*, 2001). The extent of these visible symptoms is used to describe the severity of crown rot disease, and is the predominant trait examined in disease screening experiments. Another common indicator of crown rot in the field is the premature senescence of culms (dead heads) progressing from milk to dough stages, which exhibit a whitish head containing shrivelled or no grain (Sims *et al.*, 1961; Smiley, 2019; Wallwork, 2000). Development of prematurely senescent culms becomes more frequent in crops exposed to low moisture environments during grain fill (Hollaway & Exell, 2010; Smiley, 2009; Smiley, 2019). Burgess *et al.* (2001) suggested that these culms have increased fungal colonisation and vascular disruption, a condition which has been demonstrated recently in tetraploid durum wheats (*Triticum turgidum* var. *durum*) (*Knight et al., 2017*). This investigation provided the first direct evidence of the extent of vascular colonisation occurring in prematurely senescing culms of tetraploid durum wheat, and indicated that within these culms up to 77% of vascular bundles were colonised by fungal hyphae, compared to a maximum of 14% in non-senescent culms. This was supported by visual discolouration ratings and *F. pseudograminearum* biomass estimations. A similar examination of the ‘dead head’ phenomena in hexaploid spring wheats (*Triticum aestivum*) has not previously been reported. The prematurely senescent culm condition provides opportunities to explore host and pathogen responses during extreme disease situations, and may reveal critical aspects of the crown rot disease process.

In comparison to the tetraploid durum wheats, which are considered highly susceptible to crown rot, some hexaploid spring wheats exhibit partial levels of resistance (Bovill *et al.*, 2010; Martin *et al.*, 2015; Matthews *et al.*, 2016; Wildermuth & McNamara, 1994). Genotypic variation for crown rot resistance and/or tolerance in hexaploid spring wheats provides a range of potential crown rot responses which can be exploited in breeding programs (Davies *et al.*, 2015; Kelly *et al.*, 2016). While tetraploid durum wheat and hexaploid spring wheat exhibit similar crown rot symptoms, the extent of colonisation, *F. pseudograminearum* biomass and disease progress are likely to be affected by genetic differences between the species.

The primary objective of this study was to quantify and compare disease responses, including visual discolouration, *F. pseudograminearum* biomass, vascular colonisation and host DNA content in crown rot affected hexaploid spring wheat cultivars exhibiting prematurely senescent and non-senescent culms in the field. The study was designed to complement the tetraploid durum wheat investigations of Knight *et al.* (2017). An additional objective was to examine changes in host colonisation and disease expression over time, from initial ‘dead head’ observation through to mature culms.

## Materials and methods

### Cultivar selection and culm assessment

Samples of six hexaploid spring wheat cultivars (Table 1) were collected from fields in 2014 at Narrabri, New South Wales (Plant Breeding Institute, University of Sydney), Wellcamp, Queensland (Wellcamp Field Station, Queensland Department of Agriculture and Fisheries) and Tamworth, New South Wales (Tamworth Agricultural Institute, New South Wales Department of Primary Industries).

**Table 1.** Hexaploid spring wheat cultivars assessed for visual crown rot (CR) discolouration, *Fusarium pseudograminearum* biomass, vascular colonisation and wheat DNA quantity.

| Cultivar | CR rating<sup>a</sup> | Maturity<sup>b</sup> | Location<sup>c</sup> | Inoculum<sup>d</sup> |
| --- | --- | --- | --- | --- |
| Ellison | S-VS | Mid-Fast | Narrabri | Naturalised |
| LRPB Spitfire | MS | Fast | Narrabri | Naturalised |
| Sunlin | MS-S | Mid-Fast | Narrabri | Naturalised |
| Suntop | MS-S | Mid-Fast | Narrabri | Naturalised |
| Livingston | S | Fast | Wellcamp | Naturalised/Applied |
| Sunlin T1<sup>e</sup> | MS-S | Mid-Fast | Wellcamp | Naturalised/Applied |
| Sunlin T2 | MS-S | Mid-Fast | Wellcamp | Naturalised/Applied |
| Sunlin T3 | MS-S | Mid-Fast | Wellcamp | Naturalised/Applied |
| Sunlin T4 | MS-S | Mid-Fast | Wellcamp | Naturalised/Applied |
| EGA Gregory | S | Mid-Slow | Tamworth | Natural |

<sup>a</sup>CR rating is calculated from a range of factors, including incidence of infection, dead head formation and discolouration. Rating scale: very susceptible (VS), susceptible to very susceptible (SVS), susceptible (S), moderately susceptible to susceptible (MS-S), moderately susceptible (MS), moderately resistant to moderately susceptible (MR-MS), moderately resistant (MR), resistant to moderately resistant (MR-R), and resistant (R) (Matthews *et al.* 2016).
<sup>b</sup>The maturity of selected cultivars originated from Lush *et al.* (2018) and Matthews *et al.* (2016).
<sup>c</sup>Latitude and longitude for field locations - Narrabri (-30.272931, 149.801664), Wellcamp (-27.565044, 151.864234) and Tamworth (-31.146282, 150.978124).
<sup>d</sup>The Narrabri site was under a long-term crown rot experimental trial/durum rotation over 10 years. These hexaploid spring wheats had been planted into uncultivated *F. pseudograminearum*-infected durum wheat stubble at a depth of 2.5 cm. The Wellcamp site had been maintained on a three-year rotation, consisting of one year of fallow, followed by a *F. pseudograminearum* inoculated durum wheat crop (cv. EGA Bellaroi) cultivated at maturity. The inoculated experimental trial was grown in the third year of the rotation. Further details of Wellcamp and Narrabri field site preparation, planting, inoculum production and inoculation are described by Knight & Sutherland (2015) and Knight *et al.* (2017). Tamworth samples were from a trial planted in a field with naturally occurring crown rot inoculum.
<sup>e</sup>Sunlin was collected at four time (T) points at Wellcamp. These corresponded to Zadoks growth stages 72-74 at T1, 76-78 at T2, 84-86 at T3 and 90-92 at T4 (Zadoks *et al.* 1974).

Plants from Narrabri, Wellcamp and Tamworth were selected at early milk development (growth stage 72-74, Zadoks *et al.*, 1974) from the field based on the presence of at least one prematurely senescing culm and one non-senescent, green culm at similar developmental stages. A maximum of two prematurely senescing and non-senescent culms were sampled from each plant. At Wellcamp, cv. Sunlin was also collected at three subsequent time points, each separated by 10, 9 and 16 days, respectively. These corresponded with plants at late milk development (growth stage 76-78), soft dough development (growth stage 84-86) and ripening (growth stage 90-92, with a complete loss of greenness in the plant), respectively (Zadoks *et al.*, 1974). Plants collected at early milk development were in the early stages of exhibiting prematurely senescent culm symptoms, indicated by culms still appearing green along much of their length. Prematurely senescent culm formation was indicated by a loss of greenness in the head, with this symptom continuing down into the peduncle. Frequently the flag leaf of these culms exhibited curling, with apparent dryness and a loss of greenness. Culms considered non-senescent did not exhibit these symptoms. At ripening, non-senescent and prematurely senescent culms were identified by the presence or absence of grain in the head, respectively. Culms of each plant were separated, and the leaf sheaths and roots were removed. Culm processing was as described by Knight *et al.* (2017). Briefly, each culm was sectioned into three 6 cm portions, starting at the culm base and extending up to 18 cm. In addition, 6 cm of peduncle from each culm was sampled from directly below the base of the rachis. Samples were stored at −70 °C before processing.

Visual assessment was performed on the surface area of entire 6 cm lengths of culm using a degree of discolouration scale divided into 10% increments (0, 1-10, 11-20, 21-30, 31-40, 41-50, 51-60, 61-70, 71-80, 81-90, 91-100) (Knight *et al.*, 2017). For analysis, percent discolouration was recorded as the maximum value of the range. Any brown discolouration was considered due to crown rot in the rating procedure. Genomic DNA was extracted from 15 to 20 mg of dried, ground tissue subsamples as described by Knight *et al.* (2017).

### Quantitative PCR conditions

The duplex quantitative PCR (qPCR) procedure was as described in Knight *et al.* (2012). Controls in every assay included duplicate no template controls (NTC) and genomic DNA standards (positive/negative) for both *F. pseudograminearum* and wheat. Genomic DNA standards were produced by extracting DNA from a dry weight of pure *F. pseudograminearum* mycelium (2.5 mg) or wheat seedling leaf tissue (20 mg), reflecting the weight of experimental tissue subsamples. Wheat seedling leaf tissue was collected from seven-day old seedlings of hexaploid spring wheat line 2-49, germinated on moist filter paper in a 10 cm Petri dish. Samples were dried at 50 °C overnight. DNA of standards was extracted using the Wizard Genomic DNA Extraction Kit (Promega, Sydney, NSW, Australia), following the plant DNA extraction protocol provided by the manufacturer, and quantified using a nanophotometer (Implen GmbH, München, Germany). Standards, run in triplicate, were diluted into four 10-fold serial dilutions for *F. pseudograminearum* DNA and wheat DNA. Sample qPCR values for *F. pseudograminearum* were normalised by dividing the estimated weight of *F. pseudograminearum* by the weight of extracted tissue (*F. pseudograminearum* mycelium [mg] / dry weight of tissue [g]). Sample qPCR values for wheat were normalised by dividing the estimated quantity of wheat DNA by the weight of extracted tissue (wheat DNA [ng] / dry weight of tissue [g]).

### Microscopic assessment

Transverse sections of culm tissues were taken from each sample at 1, 7 and 13 cm above the culm base. Sections were stained using the solophenyl flavine-based fluorescence method described by Knight and Sutherland (2011) and assessed for vascular colonisation using the method described by Knight *et al.* (2017). Briefly, presence and location of hyphae were determined under a fluorescence microscope (Eclipse E600, Nikon, Japan). Each large vascular bundle (those not in the sclerenchymatous hypoderm) in a section was observed and designated as colonised due to the presence of at least a single hypha in the xylem, phloem or both xylem and phloem. The depth of the parenchymatous hypoderm in each individual culm section was determined using measurements from four points in the hypoderm.

### Data analysis

Visual discolouration, *F. pseudograminearum* biomass, wheat DNA quantities, vascular colonisation (as a total and by colonisation type) and hypoderm depth were analysed for each field separately (Narrabri, Wellcamp and Tamworth). The time-course assessment of visual discolouration, *F. pseudograminearum* biomass, vascular colonisation and wheat DNA quantities was only performed at Wellcamp using cv. Sunlin. To ensure homogeneity of variance, transformations were applied to *F. pseudograminearum* biomass (log), total vascular colonisation (arcsine square root), vascular colonisation by colonisation type (log), wheat DNA quantities (log) and hypoderm depth (square root).

The analysis of each variable was performed using a linear mixed model. Model One for *F. pseudograminearum* biomass, visual discolouration, wheat DNA quantity and total vascular colonisation included culm section, cultivar, culm condition and their interactions as fixed effects. Model Two for vascular colonisation by colonisation type included the fixed effects from Model One, replacing culm section with section height, with the addition of a term to account for type of colonisation, along with the interactions between this effect and all other fixed effects in the model. Model Three for hypoderm depth included fixed effects for section height, cultivar and their interaction. The sampling factors relating to plant and culm section/section height within plant were fitted as random effects for all variables. Additionally, in Model Two, the random terms were estimated separately for each type of colonisation. These models were the same for each location except Tamworth, where all terms containing cultivar were excluded from the model as there was only one cultivar present. The same models as above were also used for the time-course assessments, the only difference being that the term for cultivar was replaced with a term to account for time.

Variance parameters were estimated using residual maximum likelihood (REML) estimation (Patterson & Thompson, 1971). Prediction of fixed effect means were generated from the model as empirical Best Linear Unbiased Estimators (eBLUEs). The analysis was performed using ASReml-R (Butler *et al.*, 2009) in the R software environment (R Core Team, 2017). Significance of fixed effects were assessed using a Wald test with a value of α=0.05.

Correlations between the different methods of disease assessment were not presented for the hexaploid spring wheats in this study due to limited variance in the data. The method of selecting the most diseased plants effectively samples one end of the distribution of disease symptoms, limiting the range of disease responses captured.

## Results

### Visual discolouration

A significant interaction was observed between culm condition (prematurely senescent or non-senescent) and culm section at Narrabri (*P* = 0.001; Table 2) and Wellcamp (*P* = 0.003; Supplementary Table S1), indicating that visual discolouration was greater in prematurely senescent culm sections than in non-senescent culm sections. Visual assessment suggested that, in general, the brown discolouration of culms became more pale from the basal 6 cm to the 12-18 cm section. There was also a significant interaction between culm condition and cultivar at Narrabri (*P* = 0.005). Cultivar Sunlin exhibited the least visual discolouration of non-senescent culms, while cv. Sunlin and cv. LRPB Spitfire exhibited the greatest visual discolouration of prematurely senescent culms. No significant effects were detected for culm condition or culm section at Tamworth. Peduncles did not exhibit visual discolouration at any location.

**Table 2.**
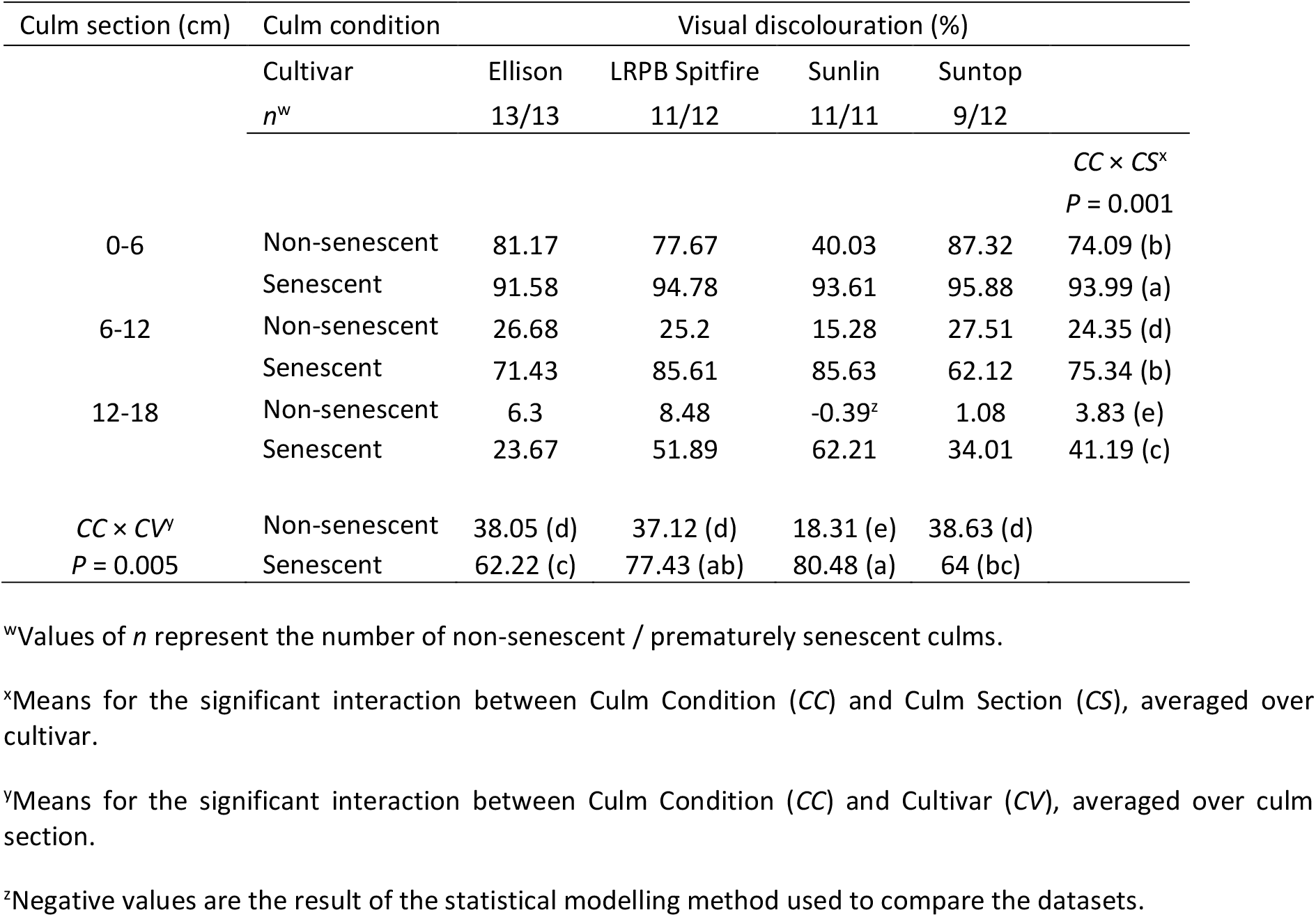
Visual discolouration (%) of each culm section of non-senescent and prematurely senescent culm conditions across hexaploid spring wheat cultivars from Narrabri. Mean discolouration ratings of each section were compared for each cultivar. Different letters indicate significant differences between groups exhibiting significant interactions.

### *Fusarium pseudograminearum* biomass

A significant effect was detected for culm condition at Narrabri (*P* < 0.001; Table 3), Wellcamp (*P* = 0.006; Supplementary Table S2) and Tamworth (*P* = 0.008; Supplementary Table S2), indicating that *F. pseudograminearum* biomass was greater in prematurely senescent culms than in non-senescent culms. No significant effect of culm section or cultivar on *F. pseudograminearum* biomass was detected at any location. No *F. pseudograminearum* biomass was detected in the peduncles.

**Table 3.**
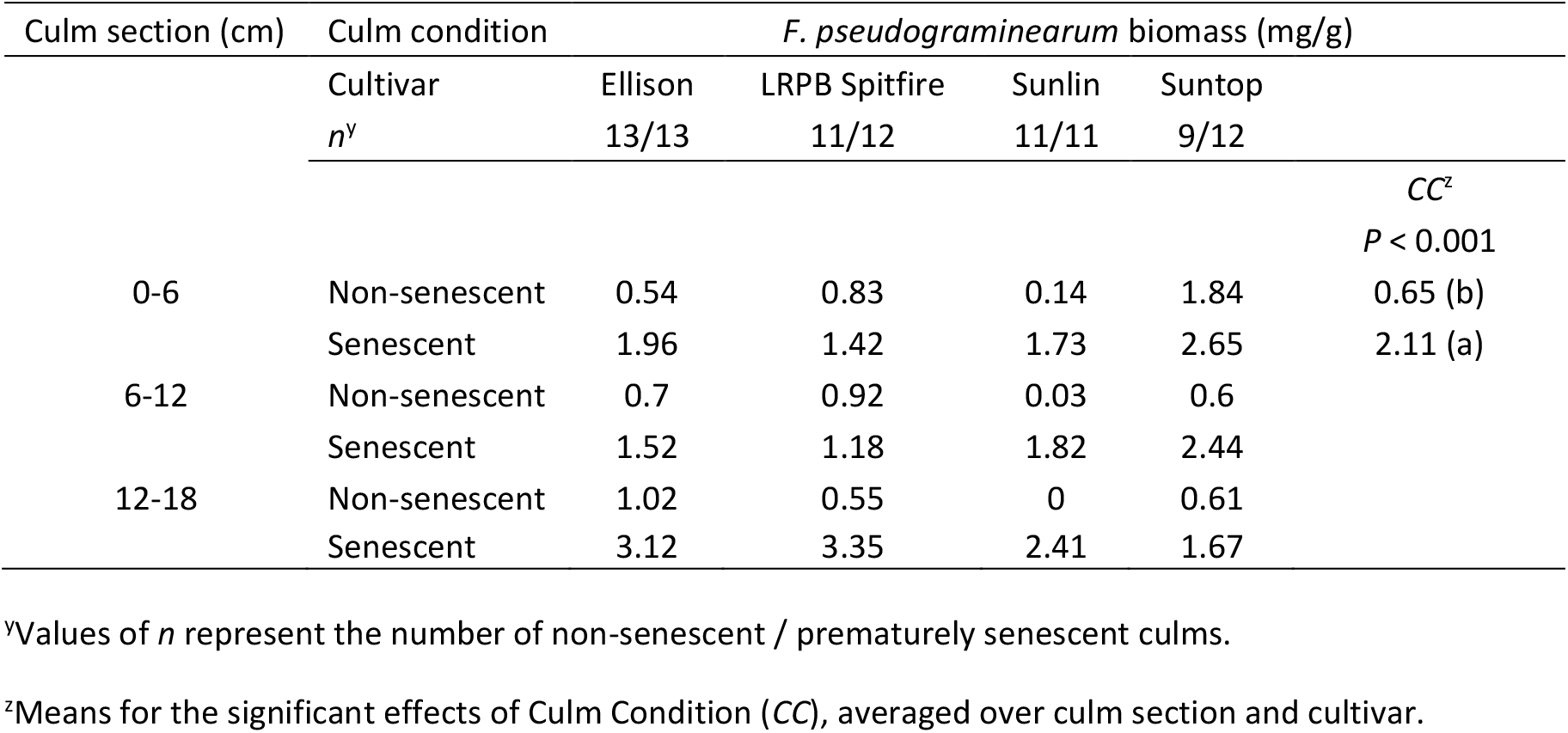
*Fusarium pseudograminearum* biomass (mg/g) of each culm section of non-senescent and prematurely senescent culm conditions across hexaploid spring wheat cultivars from Narrabri. Mean *F. pseudograminearum* biomass quantities of each section were compared among cultivars. Different letters indicate significant differences between groups exhibiting significant effects.

| Culm section (cm) | Culm condition | *F. pseudograminearum* biomass (mg/g) |  |  |  |  |
| --- | --- | --- | --- | --- | --- | --- |
|  | Cultivar | Ellison | LRPB Spitfire | Sunlin | Suntop |  |
|  | *n*<sup>y</sup> | 13/13 | 11/12 | 11/11 | 9/12 |  |
|  |  |  |  |  |  | *CC*<sup>z</sup> |
|  |  |  |  |  |  | *P* < 0.001 |
| 0-6 | Non-senescent | 0.54 | 0.83 | 0.14 | 1.84 | 0.65 (b) |
|  | Senescent | 1.96 | 1.42 | 1.73 | 2.65 | 2.11 (a) |
| 6-12 | Non-senescent | 0.7 | 0.92 | 0.03 | 0.6 |  |
|  | Senescent | 1.52 | 1.18 | 1.82 | 2.44 |  |
| 12-18 | Non-senescent | 1.02 | 0.55 | 0 | 0.61 |  |
|  | Senescent | 3.12 | 3.35 | 2.41 | 1.67 |  |

<sup>y</sup>Values of *n* represent the number of non-senescent / prematurely senescent culms.
<sup>z</sup>Means for the significant effects of Culm Condition (*CC*), averaged over culm section and cultivar.

### Vascular colonisation

Vascular colonisation of the cultivars at Narrabri exhibited a significant interaction between culm condition, section height and cultivar (*P* = 0.048; Table 4). In general, vascular colonisation was less in the non-senescent culms when compared with the prematurely senescent culms across the three culm sections in each cultivar. Analysis of vascular colonisation by colonisation type (xylem, phloem or xylem and phloem) indicated a significant interaction between culm condition, section height, cultivar and colonisation type (*P* = 0.005; Supplementary Table S3).

**Table 4.**
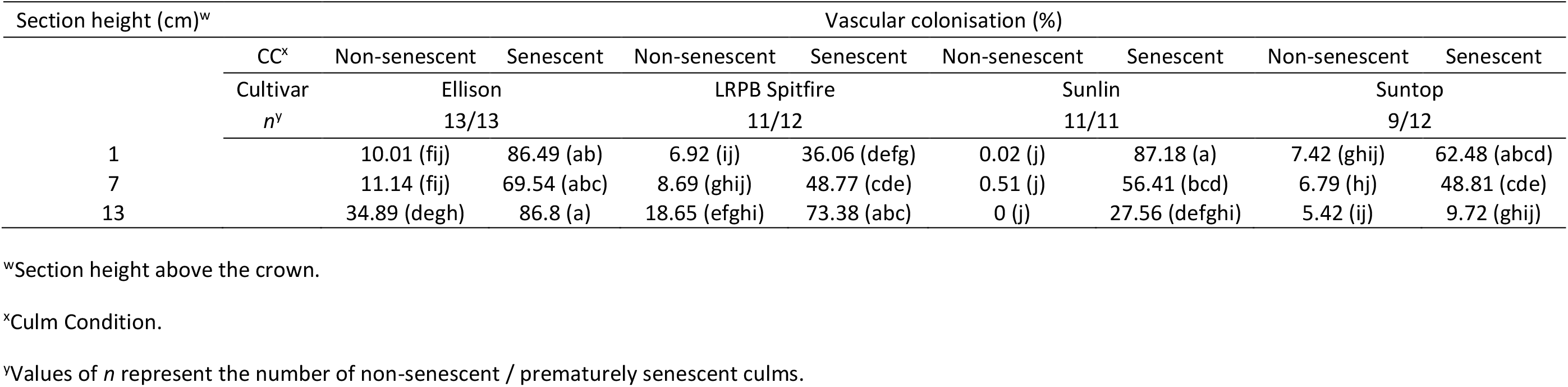
*Fusarium pseudograminearum* colonisation (%) of the vascular bundles in non-senescent and prematurely senescent culm sections of hexaploid spring wheat cultivars from Narrabri. Colonisation assessment included percentage of vascular bundles with hyphal colonisation (at least one hyphae in xylem, phloem or both xylem and phloem tissues). A significant interaction (*P* = 0.048) was observed between culm condition, section height and cultivar for the mean colonisation values. Different letters indicate significant differences between groups exhibiting significant interactions or effects.

At Wellcamp, a significant interaction occurred between culm condition and cultivar (*P* = 0.01), and between section height and cultivar (*P* < 0.001) for vascular colonisation (Supplementary Table S4). Vascular colonisation was greatest in the prematurely senescent culms of both cultivars, while the non-senescent culms of cv. Sunlin had significantly less vascular colonisation than those of cv. Livingston. Vascular colonisation at 1 cm was significantly greater than at 7 and 13 cm in cv. Livingston, while vascular colonisation at 7 cm was significantly less than at 13 cm in cv. Sunlin. Analysis of vascular colonisation by colonisation type indicated a significant interaction between culm condition, section height, cultivar and colonisation type (*P* < 0.001; Supplementary Table S3).

Vascular colonisation of cv. EGA Gregory at Tamworth exhibited a significant effect for culm condition (*P* = 0.002), with prematurely senescent culms exhibiting significantly greater vascular colonisation than non-senescent culms (Supplementary Table S4). Analysis of vascular colonisation by colonisation type indicated a significant interaction between culm condition, section height and colonisation type (*P* < 0.001; Supplementary Table S3).

While colonisation of vascular bundles was quantified, this should not be interpreted as only the vascular bundles being colonised in these sections. Consistent with previous observations (Knight *et al.*, 2017) of colonised culm tissues, hyphae were present in the epidermal cells, parenchymatous hypoderm, schlerenchymatous hypoderm and pith parenchyma.

### Wheat DNA quantities

A significant interaction was detected between culm condition, culm section and cultivar at Narrabri (*P* = 0.001) and Wellcamp (*P* = 0.002) and between culm condition and culm section at Tamworth (*P* < 0.001) for wheat DNA (Supplementary Table S5). The quantity of wheat DNA per mass of tissue decreased from the basal culm sections to the upper culm sections, while the values in the peduncle were significantly greater than in all the culm sections. No consistent pattern of differences between culm condition, culm section or cultivar was discerned at each location, however culm condition could significantly affect wheat DNA quantities. When differences were detected, they were typically prematurely senescent culm sections with lower wheat DNA content than non-senescent culm sections. However, the greatest differences occurred in the top 6 cm of peduncle tissue of cv. Ellison and cv. Sunlin, where prematurely senescent culm peduncles had significantly greater wheat DNA quantities than non-senescent peduncles.

### Hypoderm depth

Section height had a significant effect (*P* < 0.001) on parenchymatous hypoderm depth (radial thickness) across all cultivars at all three sites. The parenchymatous hypoderm depth decreased significantly from 1 cm to 7 cm and from 7 cm to 13 cm above the crown in the culms of each cultivar (Supplementary Table S6), with the exception of cv. Sunlin at Wellcamp, in which the depth of parenchymatous hypoderm was not significantly different from 7 cm to 13 cm.

### Time-course assessment of cultivar Sunlin

A significant interaction for visual discolouration was observed between culm condition and culm section across the four collection times (*P* < 0.001; Fig. 1a). This indicated significantly greater visual discolouration in prematurely senescent culms compared to non-senescent culms in the 0-6 cm and 6-12 cm sections, with no difference detected at 12-18 cm. A significant effect of time (*P* = 0.02; Fig. 1a) was also detected. When culm condition and culm section were combined by time, the visual discolouration was only significantly different between collection time 2 and 3 (63.21% and 49.61%, respectively), indicating a relatively consistent level of visual discolouration between collection times. Peduncles did not exhibit visual discolouration at any time point.

**Figure 1.**
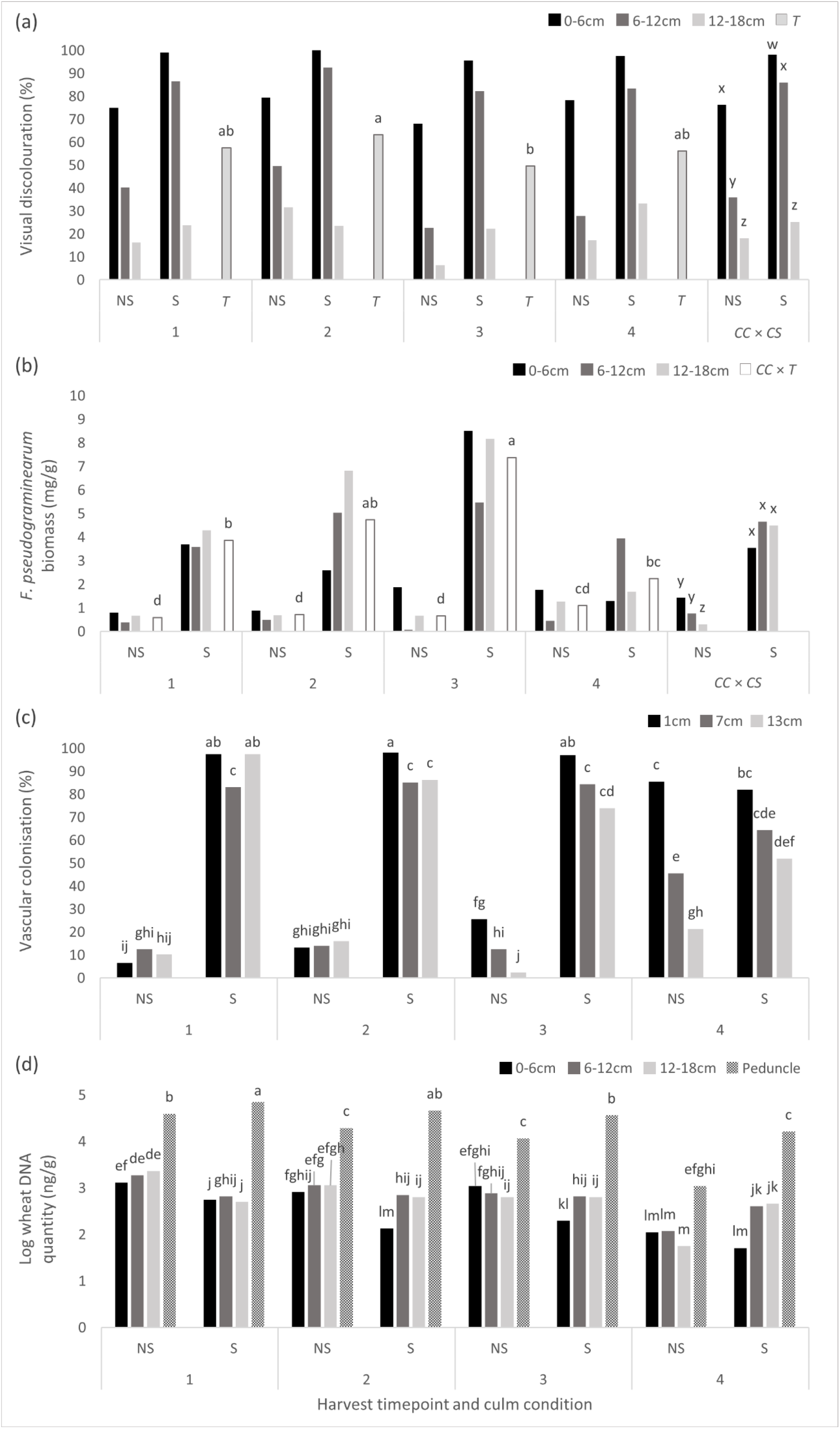
Time-course of non-senescent (NS) and prematurely senescent (S) culm conditions from hexaploid spring wheat cultivar Sunlin collected from Wellcamp at four timepoints, corresponding to early milk development (1 = Zadoks growth stage [ZGS] 72-74; *n* = 31 NS / 32 S), late milk development (2 = ZGS 76-78; *n* = 33 NS / 33 S), soft dough development (3 = ZGS 84-86; *n* = 35 NS / 37 S) and ripening (4 = ZGS 90-92; *n* = 71 NS / 9 S), respectively. Mean visual discolouration ratings (a), *Fusarium pseudograminearum* biomass quantities (b), vascular bundle colonisation values (c) and wheat DNA quantities (d) of each culm section were compared across timepoints. Culm sections included 0-6 cm, 6-12 cm and 12-18 cm from the base of the culm. Vascular colonisation was assessed at one height (1 cm, 7 cm or 13 cm, respectively) in each section. The top 6cm of peduncle tissue was also assessed, with no *F. pseudograminearum* detected. For visual discolouration, a significant interaction occurred between culm condition and culm section (*CC* × *CS*; *P* < 0.001), along with a significant effect of time (*T*; *P* = 0.02). For *F. pseudograminearum* biomass, significant interactions occurred for *CC* × *CS* (*P* < 0.001) and *CC* × *T* (*P* = 0.001). For vascular colonisation, a significant interaction occurred between culm condition, section height and time (*P* = 0.002). For wheat DNA quantities, a significant interaction occurred between culm condition, culm section and time (*P* = < 0.001). Means for significant *CC* × *CS* interactions were averaged over time, means for significant *CC* × *T* interactions were averaged over culm section, while means for significant effects of *T* were averaged over culm condition and culm section. Different letters indicate significant differences between groups exhibiting significant interactions or effects.

When each collection time was assessed for *F. pseudograminearum* biomass, a significant interaction between culm condition and culm section (*P* < 0.001) was again detected (Fig. 1b). This indicated that *F. pseudograminearum* biomass was significantly greater in prematurely senescent culms than non-senescent culms in all three culm sections. *F. pseudograminearum* biomass was significantly less in the 12-18 cm section of non-senescent culms. *F. pseudograminearum* biomass did not significantly vary between the three sections in the prematurely senescent culms.

An interaction between culm condition and time (*P* = 0.001) was also detected for *F. pseudograminearum* biomass (Fig. 1b). This demonstrated that non-senescent culms had significantly less *F. pseudograminearum* biomass than prematurely senescent culms at the first three collection times, while no significant difference was present at the fourth collection time, coinciding with plant maturity. *F. pseudograminearum* biomass in prematurely senescent culms significantly increased from the first to the third collection time and decreased from the third to the fourth collection time. *F. pseudograminearum* biomass in non-senescent culms did not significantly change between collection times.

Vascular colonisation exhibited a significant interaction between culm condition, section height and time (*P* = 0.002; Fig. 1c). Vascular colonisation was greater in the prematurely senescent culms than non-senescent culms in all three culm sections for the first three collection times. Vascular colonisation levels remained stable across collection times in the prematurely senescent culms, while the colonisation levels increased between collection times in the non-senescent culms, until reaching values similar to the prematurely senescent culm colonisation values in sections 0-6 and 6-12 cm at maturity. Analysis of vascular colonisation on the basis of each vascular tissue type (xylem, phloem or both) indicated a significant interaction between culm condition, section height, time and colonisation type (*P* < 0.001; Supplementary Table S7).

The time-course assessment of cv. Sunlin indicated a significant interaction between culm condition, culm section and time (*P* < 0.001; Fig. 1d). At collection times one and two, the non-senescent culms contained significantly greater quantities of wheat DNA than the prematurely senescent culms in sections 0-6, 6-12 and 12-18 cm. No significant differences occurred between non-senescent and prematurely senescent culms in sections 6-12 and 12-18 cm at collection time three or section 0-6 cm at collection time four. Sections 6-12 and 12-18 cm at collection time four contained significantly greater quantities of wheat DNA in the senescent culms compared to non-senescent culms. A consistent significant difference was detected in the top 6 cm of peduncle tissue, with prematurely senescent culms containing significantly greater quantities of wheat DNA than the peduncles of the non-senescent culms, at all collection times. A general decrease in wheat DNA quantities occurred from collection time one to collection time four (plant maturity) across each culm condition and culm section.

## Discussion

Prematurely senescent culms of hexaploid spring wheat exhibited increased visual discolouration, *F. pseudograminearum* biomass and vascular colonisation compared to non-senescent culms. This is the first study to comprehensively examine this colonisation in hexaploid spring wheat and follows a recent study of colonisation patterns in tetraploid durum wheats (Knight *et al.*, 2017). These comparisons begin to reveal the potential physiological impacts of *F. pseudograminearum* during cereal grain development.

Field based assessment of crown rot commonly involves assessing plants for culm browning, however prematurely senescent culms have also been used when disease levels and environmental conditions result in their production (Burgess *et al.*, 1975; Hollaway & Exell, 2010; Klein *et al.*, 1991; McKnight & Hart, 1966; Purss, 1966; Smiley, 2019). Prematurely senescent culms of both hexaploid spring and winter wheat have historically been described as having a greater level of colonisation than diseased culms classified as non-senescent, with this distinction being made based solely on the visual appearance of the wheat head and culm base (Klaasen *et al.*, 1992; Purss, 1966). Initial histopathological investigations of hexaploid spring wheat culms affected by crown rot disease reported more extensive colonisation of prematurely senescent culms, particularly in the vascular bundles (Knight & Sutherland, 2016), however this initial investigation did not provide detailed quantification of this phenomenon. The current study provides a more exhaustive investigation of the extent of colonisation in culms of commercial hexaploid spring wheat cultivars and has demonstrated that, in comparison to non-senescent culms, colonisation of both the xylem and phloem is significantly greater in prematurely senescent culms during grain fill. This reflects the findings of Knight *et al.* (2017) for tetraploid durum wheat and supports the hypothesis for prematurely senescent culm formation occurring due to compromised function of the xylem and phloem, and hyphal disruption of water transport from the root system to the head (Burgess *et al.*, 2001; Knight *et al.*, 2017).

Resistance to crown rot in hexaploid spring wheat is generally greater than that of tetraploid durum wheat, due only in part, to the presence of quantitative resistance genes in the D genome (Bovill *et al.*, 2006; Bovill *et al.*, 2010; Collard *et al.*, 2005; Collard *et al.*, 2006; Martin *et al.*, 2015; Poole *et al.*, 2012; Zheng *et al.*, 2014). Based on publicly available crown rot resistance ratings, commercial tetraploid durum wheat cultivars range from susceptible to very susceptible, while some recently released commercial hexaploid spring wheats rate as moderately resistant to moderately susceptible (Matthews *et al.*, 2016). The current study was biased towards more susceptible cultivars (Table 1), which readily produced prematurely senescent culms under disease conditions in the field. Despite this, cultivar effects were observed for visual discolouration and vascular colonisation. These differences between cultivars were variable by culm section, and did not separate the cultivars by crown rot resistance rating. Indeed, variation in disease levels between individual culms collected from single plants may be as significant as the variation observed among different genetic backgrounds, demonstrating the on-going challenge of describing significant differentiation of crown rot reactions in the field.

The three disease assessment procedures applied in this investigation have provided insights into different aspects of the disease system. Visual discolouration provides information on the development and accumulation of visible disease symptoms up to the time of collection and is analogous to a measure of the area under a disease progress curve (AUDPC). In contrast, values of *F. pseudograminearum* biomass and vascular colonisation have potential to vary in response to disease state and age of the host tissue at the time of collection. Visual discolouration cannot differentiate between cultivar responses once 100% discolouration occurs, and is hypothesised to be affected by a decrease in the depth of parenchymatous hypoderm at increasing heights above the culm base along hexaploid spring and tetraploid durum wheat culms (Knight & Sutherland, 2016; Knight *et al.*, 2017). Comparing visual discolouration ratings at different positions in a culm may be impacted by structural differences, as well as by potential variation in structure, such as parenchymatous hypoderm depth, among cultivars. Further investigation of the relationship between hypoderm depth and visual discolouration across a wider range of cultivars is required. In contrast to visual discolouration, no significant effect of culm section was detected for *F. pseudograminearum* biomass, which indicates similar colonisation levels along the basal 18 cm of culm tissues at early milk development. While no colonisation was detected in peduncle tissues, further investigation of the extent of colonisation along entire culms may be informative. Quantitative estimates of disease reactions such as fungal biomass, colonisation or relative yield loss are necessary. The effects of the crown rot resistance rating and biomass of postharvest stubble on saprophytic growth and inoculum levels carried over between seasons should also be considered (Summerell *et al.*, 1989).

Observations of crown rot disease have predominantly focussed on disease conditions directly associated with the fungus, however a greater understanding of the plant physiological responses to disease have been highlighted as essential for crown rot resistance research (Knight et al., 2017). The first report on changes in host DNA quantities in relation to disease have been provided in this study, where a range of significant responses across cultivars, geographic regions and time were observed. Significant differences between culm conditions varied, and in general the prematurely senescent culms contained less wheat DNA in the basal culm sections than the non-senescent culms, presumably due to fungal colonisation and resultant host cell death. In contrast, the greatest differences observed between culm conditions occurred in the peduncle, where no *F. pseudograminearum* was detected, and prematurely senescent culms contained significantly greater quantities of wheat DNA than non-senescent culms in two cultivars (Ellison and Sunlin). The trend of significantly more wheat DNA in peduncles of prematurely senescent culms persisted into maturity for cv. Sunlin, along with a consistent decrease in wheat DNA quantities along the culms during the transition from the milk development stage into maturity. It is hypothesised that the rapid senescence occurring in prematurely senescent culms may limit typical programmed cell death processes in the peduncle, resulting in significantly greater quantities of wheat DNA remaining intact. Hence, while the wheat DNA in peduncles of non-senescent culms undergoes programmed breakdown reflective of typical maturation processes, in contrast, the peduncles of the prematurely senescent culms experienced reduced loss of DNA, related to the earlier timing of the premature senescence initiation event which is accompanied by a rapid and much earlier drying of the tissue.

The diverse genetic backgrounds of the cultivars, particularly with respect to maturity, impeded informative comparisons of the wheat DNA related observations. It would be ideal to examine these effects using resources such as near isogenic lines or lines with similar maturation times from doubled haploid populations which exhibit differences in disease reactions. These observations confirm the complexity of interactions occurring between the host and fungus during plant and disease development, while recent studies suggest the involvement of secondary metabolites (Powell et al., 2017; Sørensen et al., 2018). Further investigation into the effects of disease on physiological processes such as plant maturation, phloem transport, xylem function, transpiration rates and photosynthesis is required.

The variation in wheat DNA quantities between host tissues, cultivars and time points also suggests that the use of this measure as a normalising factor for estimations of pathogen DNA content in infected tissues is not ideal. Knight et al. (2012) described similar variation in cereal seedling tissues, and supported the use of plant tissue dry weights as an alternative normalising factor.

Timing of tissue collection may have a strong influence on crown rot disease measurement, along with individual cultivar growth patterns, growth stage and environmental variation (Hogg *et al.*, 2007; Knight *et al.*, 2012; Knight & Sutherland, 2015; Knight & Sutherland, 2016; Knight *et al.*, 2017; Percy *et al.*, 2012; Smiley, 2019; Stephens *et al.*, 2008; Summerell *et al.*, 1989). The current study included a time-course examination of the impact of plant maturation on crown rot disease responses. While assessment was only performed in one year, the findings support those of Knight and Sutherland (2015), in which a plateau or reduction in *F. pseudograminearum* biomass was observed in two hexaploid spring wheat genotypes at maturity. The current study demonstrated that within cv. Sunlin, visual discolouration remained constant up to maturity, while *F. pseudograminearum* biomass peaked in prematurely senescent culms at soft dough development, followed by a decrease at crop maturity. Non-senescent culms, however, never reached the same peak of *F. pseudograminearum* biomass observed in the prematurely senescent culms. At crop maturity, *F. pseudograminearum* biomass in both culm conditions was similar. Vascular colonisation increased in the non-senescent culms up to maturity, reflecting the results of Knight and Sutherland (2016) for the susceptible cv. Puseas, while colonisation in the prematurely senescent culms was consistent over the sampling period. The increase in vascular colonisation at maturity in the non-senescent culms may be due to increased growth of *F. pseudograminearum* during the moisture stress occurring during this process (Beddis & Burgess, 1992; Clement & Parry, 1998). This increased colonisation is hypothesised to have occurred earlier in the prematurely senescent culms. The temporally distinct patterns of colonisation observed between non-senescent and prematurely senescent culms may suggest variation in the time of initial infection among culms of the same plant. Further investigation of the timing of initial infection of each culm, and the factors associated with differential responses within plants, is required.

## Acknowledgments

The authors thank Dr. Philip Davies (Plant Breeding Institute, University of Sydney) and Dr Steven Simpfendorfer (Tamworth Agricultural Institute, NSW DPI) for providing field samples. Financial support for this project was provided by the University of Southern Queensland Strategic Research Fund.

## Notes

### Competing Interest Statement

The authors have declared no competing interest.

