## Supplementary Table for "Disease responses of hexaploid spring wheat (*Triticum aestivum*) culms exhibiting premature senescence (dead heads) associated with *Fusarium pseudograminearum* crown rot"

**Supplementary Table S1** Visual discolouration (%) of each culm section of non-senescent and prematurely senescent culm conditions across hexaploid spring wheat cultivars from Wellcamp and Tamworth. Mean discolouration ratings of each section were compared for each cultivar within each location. Different letters indicate significant differences between groups exhibiting significant interactions. Significant interactions between culm condition and culm section were not observed for cultivar EGA Gregory at Tamworth.

| Location | Culm section (cm) | Culm condition | Visual discolouration (%) | | | |
|---|---|---|---|---|---|---|
| Wellcamp | | Cultivar | Livingston | Sunlin | - | |
| | | $n^w$ | 28/28 | 31/32 | - | |
| | | | | | | *CC* × *CS*[x] <br> *P* = 0.003 |
| | 0-6 | Non-senescent | 75.04 | 73.27 | - | 74.15 (b) |
| | | Senescent | 92.47 | 99.95 | - | 96.74 (a) |
| | 6-12 | Non-senescent | 32.18 | 38.33 | - | 35.26 (c) |
| | | Senescent | 64 | 89.13 | - | 78.36 (b) |
| | 12-18 | Non-senescent | 6.88 | 15.24 | - | 11.06 (d) |
| | | Senescent | 6.88 | 26.45 | - | 18.07 (d) |
| Tamworth | | Cultivar | - | - | EGA Gregory | - |
| | | *n* | - | - | 6/7 | - |
| | 0-6 | Non-senescent | - | - | 88.17 | - |
| | | Senescent | - | - | 94.79 | - |
| | 6-12 | Non-senescent | - | - | 62.73 | - |
| | | Senescent | - | - | 89.09 | - |
| | 12-18 | Non-senescent | - | - | 67.19 | - |
| | | Senescent | - | - | 54.48 | - |

[w]Values of *n* represent the number of non-senescent / prematurely senescent culms.
[x]Means for the significant interaction between Culm Condition (*CC*) and Culm Section (*CS*), averaged over cultivar.

**Supplementary Table S2** *Fusarium pseudograminearum* biomass (mg/g) of each culm section of non-senescent and prematurely senescent culm conditions across hexaploid spring wheat cultivars from Wellcamp and Tamworth. Mean *F. pseudograminearum* biomass quantities of each section were compared among cultivars within each location. Different letters indicate significant differences between groups exhibiting significant effects.

| Location | Culm section (cm) | Culm condition | *F. pseudograminearum* biomass (mg/g) | | | |
|---|---|---|---|---|---|---|
| Wellcamp | | Cultivar | Livingston | Sunlin | - | |
| | | *n*[y] | 28/28 | 31/32 | - | |
| | | | | | | *CC*[z] |
| | | | | | | $P$ = 0.006 |
| | 0-6 | Non-senescent | 1.68 | 0.76 | - | 0.76 (b) |
| | | Senescent | 8.14 | 4.42 | - | 3.77 (a) |
| | 6-12 | Non-senescent | 0.87 | 0.6 | - | |
| | | Senescent | 1.46 | 3.64 | - | |
| | 12-18 | Non-senescent | 0.33 | 0.32 | - | |
| | | Senescent | 1.99 | 2.98 | - | |
| Tamworth | | Cultivar | - | - | EGA Gregory | |
| | | *n* | - | - | 6/7 | |
| | | | | | | *CC* |
| | | | | | | $P$ = 0.008 |
| | 0-6 | Non-senescent | - | - | 4.73 | 2.27 (b) |
| | | Senescent | - | - | 3.92 | 8.75 (a) |
| | 6-12 | Non-senescent | - | - | 0.34 | |
| | | Senescent | - | - | 3.47 | |
| | 12-18 | Non-senescent | - | - | 1.75 | |
| | | Senescent | - | - | 18.86 | |

[y]Values of *n* represent the number of non-senescent / prematurely senescent culms.
[z]Means for the significant effects of Culm Condition (*CC*), averaged over culm section and cultivar.

**Supplementary Table S3** *Fusarium pseudograminearum* colonisation of the xylem, phloem or xylem and phloem of vascular bundles in non-senescent and prematurely senescent culm sections of spring wheat cultivars from three locations. Colonisation assessment included percentage of vascular bundles with hyphal colonisation (at least one hypha in xylem, phloem or both xylem and phloem tissues). Mean colonisation (%) values of each section were compared for significant interactions between culm condition, section height, cultivars and colonisation type. Different letters indicate significant differences between groups exhibiting significant interactions. Phloem data were excluded from the analysis due to frequent zero values.

| Location | Colonisation type | Section height (cm)[a] | | Colonisation (%) | | | | | | | |
|---|---|---|---|---|---|---|---|---|---|---|---|
| | | | CC[b] | Non-senescent | Senescent | Non-senescent | Senescent | Non-senescent | Senescent | Non-senescent | Senescent |
| | | | Cultivar | Ellison | | LRPB Spitfire | | Sunlin | | Suntop | |
| | | | *n*[c] | 13/13 | | 11/12 | | 11/11 | | 9/12 | |
| Narrabri | No colonisation | 1 | | 90.43 (abce) | 13.15 (mnopqrstuvwxyzAB) | 93.13 (abcd) | 63.86 (efghij) | 99.98 (a) | 12.97 (mnopqrstuvwxyzABC) | 92.78 (abcde) | 36.66 (hijklmno) |
| | | 7 | | 88.62 (abcde) | 30.35 (jlmnop) | 91.53 (abcde) | 51.1 (fghijk) | 99.43 (a) | 44.04 (ghijkl) | 93.11 (abcde) | 51.64 (fghijk) |
| | | 13 | | 66.31 (dfghi) | 12.82 (mnopqrstuvwxyzAB) | 81.32 (bcdef) | 26.36 (klmnopqrs) | 100 (a) | 72.42 (bcdefg) | 94.74 (ab) | 90.53 (abcde) |
| | Xylem and phloem | 1 | | 3.4 (tuvwxyzABCDEF) | 68.74 (bcdefgh) | 2.85 (tuvwxyzABCDEF) | 17.5 (lmnopqrstuvwxyz) | 0 (DE) | 67.31 (cdefghi) | 0.42 (ADEF) | 35.24 (ijklmo) |
| | | 7 | | 1.45 (zACDEF) | 29.76 (klmnop) | 2.66 (tuvwxyzABCDEF) | 27.27 (klmnopqr) | 0 (DE) | 24.96 (klmnopqrs) | 1.84 (wyzACDEF) | 20.22 (klmnopqrstuv) |
| | | 13 | | 17.38 (lmnopqrstuvwxy) | 65.67 (efghi) | 8.55 (oqrstuvwxyzABCD) | 28.15 (klmnopq) | 0 (DE) | 2.56 (tuvwxyzABCDEF) | 1.09 (wACDEF) | 4.9 (ruvwxyzABCDEF) |
| | Xylem | 1 | | 3.48 (uvwxzABCDEF) | 11.66 (pqrstuvwxyzA) | 2.89 (xABCDEF) | 5.63 (tuvwxyzABCD) | 0.02 (EF) | 12 (npqrstuvwxyzA) | 7.93 (qrstuvwxyzABC) | 15.96 (mnopqrsty) |
| | | 7 | | 8.2 (qrstuvwxyzAB) | 30.69 (klm) | 4.19 (tuvwxyzABCDF) | 9.85 (pqrstuvwxyzAB) | 0.6 (CDEF) | 21.41 (lmnopqr) | 3.46 (vxABCDEF) | 14.47 (mnopqrstuwyz) |
| | | 13 | | 6.88 (stuvwxyzABCD) | 10.13 (pqrstuvwxyzAB) | 4.49 (tuvwxyzABCDF) | 29.16 (klmo) | 0 (E) | 20.91 (lmnopqr) | 3.49 (uvwxzABCDEF) | 1.94 (BCDEF) |
| | Phloem[d] | 1 | | 2.69 | 6.45 | 1.13 | 13.01 | 0 | 7.72 | 0 | 12.14 |
| | | 7 | | 1.73 | 9.2 | 1.62 | 11.78 | 0 | 9.59 | 1.59 | 13.67 |
| | | 13 | | 9.43 | 11.38 | 5.64 | 16.33 | 0 | 4.11 | 0.68 | 2.63 |
| | *CC:SH:CV:Col*[e] | *P* = 0.005 | | | | | | | | | |
| | | | Cultivar | Livingston | | Sunlin | | - | | - | |
| | | | *n* | 28/28 | | 31/32 | | - | | - | |
| Wellcamp | No colonisation | 1 | | 71.33 (c) | 3.37 (iklm) | 96.51 (a) | 1.04 (lm) | - | - | - | - |
| | | 7 | | 77.39 (bc) | 12.96 (fghj) | 90.97 (ab) | 16.42 (fg) | - | - | - | - |
| | | 13 | | 94.48 (a) | 10.99 (fghij) | 93.17 (a) | 0.96 (lm) | - | - | - | - |
| | Xylem and phloem | 1 | | 13.87 (fghi) | 88.64 (ab) | 1.57 (kl) | 87.96 (ab) | - | - | - | - |
| | | 7 | | 4.97 (ghijklm) | 38.59 (e) | 3.24 (jklm) | 40.18 (de) | - | - | - | - |
| | | 13 | | 3.23 (jklm) | 58.86 (cd) | 2.75 (jklm) | 88.53 (ab) | - | - | - | - |
| | Xylem | 1 | | 8.81 (ghij) | 4.73 (hijkm) | 0.75 (lm) | 8.38 (ghij) | - | - | - | - |
| | | 7 | | 10.47 (fghij) | 38.89 (e) | 2.37 (lm) | 34.07 (e) | - | - | - | - |
| | | 13 | | 0.66 (l) | 18.38 (f) | 0.91 (lm) | 7.9 (ghijk) | - | - | - | - |

| | | | | | | | | | | |
|---|---|---|---|---|---|---|---|---|---|---|
| | | 1 | 5.99 | 3.26 | 1.17 | 2.62 | - | - | - | - |
| | Phloem | 7 | 7.17 | 9.56 | 3.42 | 9.33 | - | - | - | - |
| | | 13 | 1.63 | 11.77 | 3.17 | 2.61 | - | - | - | - |
| | *CC:SH:CV:Col* | *P* < 0.001 | | | | | | | | |
| | | Cultivar | EGA Gregory | | - | | - | | - | |
| | | *n* | 6/7 | | - | | - | | - | |
| Tamworth | | 1 | 64.37 (abcde) | 0.59 (i) | - | - | - | - | - | - |
| | No colonisation | 7 | 73.74 (abc) | 11.55 (fghi) | - | - | - | - | - | - |
| | | 13 | 79.18 (ab) | 25.85 (efgh) | - | - | - | - | - | - |
| | | 1 | 5.76 (ghi) | 96.47 (a) | - | - | - | - | - | - |
| | Xylem and phloem | 7 | 11.05 (fghi) | 62.75 (bce) | - | - | - | - | - | - |
| | | 13 | 8.07 (ghi) | 25.95 (dfgh) | - | - | - | - | - | - |
| | | 1 | 25.43 (dfg) | 1.81 (i) | - | - | - | - | - | - |
| | Xylem | 7 | 10.27 (ghi) | 24.55 (fg) | - | - | - | - | - | - |
| | | 13 | 5.08 (hi) | 36.96 (cdef) | - | - | - | - | - | - |
| | | 1 | 4.44 | 1.13 | - | - | - | - | - | - |
| | Phloem | 7 | 4.94 | 1.15 | - | - | - | - | - | - |
| | | 13 | 7.67 | 11.24 | - | - | - | - | - | - |
| | *CC:SH:Col*[f] | *P* < 0.001 | | | | | | | | |

[a]Section height above the crown.
[b]Culm Condition.
[c]Values of *n* represent the number of non-senescent / prematurely senescent culms.
[d]Phloem colonisation was omitted from analysis due to a high number of zero scores.
[e]Significant interaction between Culm Condition (*CC*), Section Height (*SH*), Cultivar (*CV*) and Colonisation Type (*Col*)
[f]Significant interaction between Culm Condition (*CC*), Section Height (*SH*) and Colonisation Type (*Col*)

**Supplementary Table S4** *Fusarium pseudograminearum* colonisation (%) of the xylem and phloem of vascular bundles in non-senescent and prematurely senescent culm sections of hexaploid spring wheat cultivars from Wellcamp and Tamworth. Colonisation assessment included percentage of vascular bundles with hyphal colonisation (at least one hyphae in xylem, phloem or both xylem and phloem tissues). Mean colonisation values of each section were compared for significant interactions between culm condition, section height and cultivars. Different letters indicate significant differences between groups exhibiting significant interactions or effects.

| Location | Section height (cm)[u] | Vascular colonisation (%) | | | | | | | | |
|---|---|---|---|---|---|---|---|---|---|---|
| | | CC[v] | Non-senescent | Senescent | Non-senescent | Senescent | Non-senescent | Senescent | *SH × CV*[w] | |
| Wellcamp | | Cultivar | Livingston | | Sunlin | | - | | Livingston | Sunlin |
| | | *n*[x] | 28/28 | | 31/32 | | - | | 56 | 63 |
| | | | | | | | - | | $P$ = <0.001 | |
| | 1 | | 28.68 | 96.65 | 3.45 | 98.96 | - | - | 68.51 (a) | 54.29 (abc) |
| | 7 | | 22.71 | 86.92 | 9.04 | 83.48 | - | - | 56.21 (bc) | 44.33 (c) |
| | 13 | | 5.54 | 89.01 | 6.97 | 99.04 | - | - | 45.01 (bc) | 58.46 (ab) |
| *CC × CV*[y] | $P$ = 0.01 | | 17.62 (b) | 91.41 (a) | 6.27 (c) | 95.81 (a) | | | | |
| Tamworth | | Cultivar | - | | - | | EGA Gregory | | - | |
| | | *n* | - | | - | | 6/7 | | - | |
| | 1 | | - | - | - | - | 35.93 | 99.4 | - | - |
| | 7 | | - | - | - | - | 26.51 | 88.29 | - | - |
| | 13 | | - | - | - | - | 21.05 | 74.34 | - | - |
| *CC*[z] | $P$ = 0.002 | | | | | | 27.66 (b) | 90.15 (a) | | |

[u]Section height above the crown.
[v]Culm Condition.
[w]Means for the significant interaction of Section Height (*SH*) and Cultivar (*CV*), averaged over culm condition.
[x]Values of *n* represent the number of non-senescent / prematurely senescent culms.
[y]Means for the significant interaction of Culm Condition (*CS*) and Cultivar (*CV*), averaged over section height.
[z]Means for the significant effect of Culm Condition (*CC*), averaged over section height.

**Supplementary Table S5** Wheat DNA quantity (ng/g) of each culm section of non-senescent and prematurely senescent culm conditions across hexaploid spring wheat cultivars from three locations. Mean wheat DNA quantities of each section were compared for each cultivar within each location. Different letters indicate significant differences between groups exhibiting significant interactions.

| Location | Culm section (cm) | Culm condition | Wheat DNA (ng/g) | | | |
|---|---|---|---|---|---|---|
| Narrabri | | Cultivar | Ellison | LRPB Spitfire | Sunlin | Suntop |
| | | $n^w$ | 13/13 | 11/12 | 11/11 | 9/12 |
| | 0-6 | Non-senescent | 786 (fghijklm) | 904 (efghij) | 1516 (efg) | 2023 (e) |
| | | Senescent | 542 (hijklmno) | 975 (efghi) | 663 (hijklmn) | 861 (fghijk) |
| | 6-12 | Non-senescent | 364 (jklmno) | 793 (ghijkl) | 434 (ijklmno) | 535 (hijklmno) |
| | | Senescent | 375 (jklmno) | 329 (lmno) | 233 (o) | 813 (fghijkl) |
| | 12-18 | Non-senescent | 343 (klmno) | 363 (klmno) | 339 (klmno) | 761 (fghijklmn) |
| | | Senescent | 339 (klno) | 1731 (ef) | 589 (hijklmno) | 1123 (efgh) |
| | Peduncle[x] | Non-senescent | 15026 (d) | 46126 (ab) | 22684 (bcd) | 28079 (abcd) |
| | | Senescent | 53302 (a) | 36685 (abc) | 48444 (a) | 18321 (cd) |
| | *CC × CS × CV*[y] | *P* = 0.001 | | | | |
| Wellcamp | | Cultivar | Livingston | Sunlin | - | - |
| | | *n* | 28/28 | 31/32 | - | - |
| | 0-6 | Non-senescent | 863 (fgh) | 1390 (ef) | - | - |
| | | Senescent | 537 (hi) | 610 (ghi) | - | - |
| | 6-12 | Non-senescent | 1087 (fg) | 1920 (de) | - | - |
| | | Senescent | 720 (gh) | 713 (ghi) | - | - |
| | 12-18 | Non-senescent | 2707 (d) | 2381 (d) | - | - |
| | | Senescent | 343 (i) | 576 (ghi) | - | - |
| | Peduncle | Non-senescent | 17524 (c) | 39831 (b) | - | - |
| | | Senescent | 13335 (c) | 71664 (a) | - | - |
| | *CC × CS × CV*[y] | *P* = 0.002 | | | | |
| Tamworth | | Cultivar | EGA Gregory | - | - | - |
| | | *n* | 6/7 | - | - | - |
| | 0-6 | Non-senescent | 2025 (b) | - | - | - |
| | | Senescent | 83 (d) | - | - | - |
| | 6-12 | Non-senescent | 261 (cd) | - | - | - |
| | | Senescent | 432 (cd) | - | - | - |
| | 12-18 | Non-senescent | 646 (bc) | - | - | - |
| | | Senescent | 763 (bc) | - | - | - |
| | Peduncle | Non-senescent | 20268 (a) | - | - | - |
| | | Senescent | 40815 (a) | - | - | - |
| | *CC × CS*[z] | *P* < 0.001 | | | | |

[w]Values of *n* represent the number of non-senescent / prematurely senescent culms.
[x]Top 6 cm of the peduncle.
[y]Significant interactions between Culm Condition (*CC*), Culm Section (*CS*) and Cultivar (*CV*).
[z]Significant interactions between Culm Condition (*CC*) and Culm Section (*CS*).

**Supplementary Table S6** Hypoderm depth at three positions in spring wheat culms from three locations. Mean hypoderm depth (µm) was compared between each position in a culm across cultivars in each location. Different letters indicate significant differences between groups exhibiting significant interactions or effects.

| Location | Section height (cm)[w] | Hypoderm depth (µm) | | | | |
|---|---|---|---|---|---|---|
| | Cultivar | Ellison | LRPB Spitfire | Sunlin | Suntop | |
| | *n*[x] | 26 | 23 | 22 | 21 | |
| | | | | | | *SH*[y] |
| | | | | | | *P* < 0.001 |
| Narrabri | 1 | 59.99 | 61.36 | 59.31 | 58.79 | 59.68 (a) |
| | 7 | 2.12 | 6.67 | 10.1 | 11.26 | 7.1 (b) |
| | 13 | 0.28 | 1.38 | 1.06 | 1.15 | 0.88 (c) |
| | | | | | | - |
| | Cultivar | Livingston | Sunlin | - | - | - |
| | *n* | 56 | 63 | - | - | - |
| Wellcamp | 1 | 48.81 (a) | 57.68 (a) | - | - | - |
| | 7 | 7.52 (b) | 0.7 (c) | - | - | - |
| | 13 | 1.18 (c) | 0.28 (c) | - | - | - |
| | | | | | | - |
| | *SH:CV*[z] | *P* < 0.001 | | | | - |
| | Cultivar | EGA Gregory | - | - | - | - |
| | *n* | 13 | - | - | - | - |
| Tamworth | 1 | 79.47 (a) | - | - | - | - |
| | 7 | 23.18 (b) | - | - | - | - |
| | 13 | 5.35 (c) | - | - | - | - |
| | | | | | | - |
| | *SH* | *P* < 0.001 | | | | - |

[w]Section height above the crown.
[x]Values of *n* represent the number of culms assessed.
[y]Means for the significant effects of Section Height (*SH*), averaged over cultivar.
[z]Significant interaction between Section Height (*SH*), and Cultivar (*CV*).

**Supplementary Table S7** *Fusarium pseudograminearum* colonisation of the xylem, phloem or xylem and phloem of vascular bundles in non-senescent and prematurely senescent culm sections collected at four timepoints, corresponding to early milk development (Zadoks growth stage 72-74), late milk development (Zadoks growth stage 76-78), soft dough development (Zadoks growth stage 84-86) and ripening (Zadoks growth stage 90-92), respectively. Mean colonisation (%) values of each section were compared across each timepoint. Different letters indicate significant differences between groups exhibiting significant interactions. Phloem data were excluded from the analysis due to frequent zero values.

| Colonisation type | Section height (cm)[a] | | Colonisation (%) | | | | | | | |
|---|---|---|---|---|---|---|---|---|---|---|
| | | CC[b] | Non-senescent | Senescent | Non-senescent | Senescent | Non-senescent | Senescent | Non-senescent | Senescent |
| | | Time | 1 | | 2 | | 3 | | 4 | |
| | | *n*[c] | 31/32 | | 33/33 | | 35/37 | | 71/9 | |
| No colonisation | 1 | | 96.32 (ab) | 1.07 (HIJK) | 89.89 (bcd) | 0.67 (IJK) | 77.61 (de) | 1.17 (GIJK) | 12.97 (ruvwxyA) | 16.67 (nortuvwxyzABCDEF) |
| | 7 | | 90.81 (bcd) | 16.29 (ortuvwxy) | 88.79 (bcd) | 13.54 (rtuvwxyAB) | 91.07 (bc) | 15.16 (rstuvwxyA) | 54.9 (fg) | 35.2 (fghijklmnopqrst) |
| | 13 | | 93.3 (ab) | 1.01 (GIJK) | 88 (bcd) | 12.01 (ruvwxyzABC) | 99.38 (a) | 25.16 (klmnopqrstu) | 81.53 (cd) | 49.09 (efghijklm) |
| Xylem and phloem | 1 | | 1.6 (EFGHIJK) | 87.62 (bcd) | 3.24 (zCDEFGHIJK) | 90.53 (bcd) | 8.13 (vwxyzABCDEGH) | 93.27 (ab) | 51.62 (fg) | 50.35 (fghijk) |
| | 7 | | 3.31 (zBCDEFGHIJK) | 40.35 (fghijkmn) | 1.33 (GIJK) | 46.56 (fghj) | 1.65 (FIJK) | 44.8 (fghi) | 12.6 (ruvwxyA) | 25.11 (hijklmnopqrstuvwx) |
| | 13 | | 2.65 (zCDEFGHIJK) | 88.62 (bcd) | 4.86 (yzABCDEFGHIK) | 58.22 (f) | 0.3 (IJ) | 30.31 (jklmnopqs) | 4.41 (zBCDEFGHK) | 6.02 (uzABCDEFGHIJK) |
| Xylem | 1 | | 1 (IJK) | 8.84 (vwxyzABCD) | 3.83 (zCDEFGHIK) | 6.75 (yzABCDEFGH) | 7.53 (xyzABCDEFH) | 3.34 (CDEFGHIK) | 24.8 (lopqst) | 21.65 (mnopqrstuvw) |
| | 7 | | 2.45 (EFGHIJK) | 33.19 (hijklmnq) | 7.44 (xyzABCDEFGH) | 29.67 (klmnopq) | 5.37 (zABCDEFGHK) | 30.82 (iklmnoq) | 20.45 (prstu) | 18.28 (oqrstuvwxy) |
| | 13 | | 1.03 (IJK) | 7.71 (wxyzABCDEF) | 3.18 (DEFGHIK) | 17.64 (rstuv) | 0.17 (J) | 26.59 (klmnopqst) | 8.8 (wxyzAB) | 35.71 (ghijklmnopq) |
| Phloem[d] | 1 | | 1.08 | 2.47 | 3.04 | 2.05 | 6.73 | 2.22 | 10.61 | 11.33 |
| | 7 | | 3.44 | 10.17 | 2.44 | 10.23 | 1.91 | 9.22 | 12.05 | 21.41 |
| | 13 | | 3.02 | 2.66 | 3.96 | 12.13 | 0.15 | 17.94 | 5.26 | 9.18 |
| *CC:SH:T:Col*[e] | $P < 0.001$ | | | | | | | | | |

[a]Section height above the crown.
[b]Culm Condition.
[c]Values of *n* represent the number of non-senescent / prematurely senescent culms.
[d]Phloem colonisation was omitted from analysis due to a high number of zero scores.
[e]Significant interaction between Culm Condition (*CC*), Section Height (*SH*), Time (*T*) and Colonisation Type (Col).
